# Vape Aerosol Exposure Decreases Voluntary Exercise in Mice

**DOI:** 10.64898/2026.10.12.758346

**Authors:** Dylan Dawood, Clarissa Savko, Sama Mikhail, Pria Bose, Morgan K. Wright, Carolina Esquer, Barbara Bailey, Joy A. Phillips, Mark Sussman

## Abstract

Electronic cigarette use (vaping) is generally accepted as a less harmful alternative to traditional cigarette smoking. However, so called “never smokers” including young adults and teenagers are often recruited through targeted advertising and appealing vape juice flavors. Any fitness cost of vaping matters for young people who exercise or play sports. We therefore tracked running distance in a mouse model of voluntary exercise to assess the effect of vaping. Specifically, we asked whether vaping decreases exercise performance, and whether there is progressive change over time. Experimental groups of mice subjected to whole-body inhalation exposure and corresponding non-exposed controls were housed in cages with running wheels. Daily wheel laps were recorded over the 8-week exposure period and analyzed using hierarchical generalized additive mixed models. Vape-exposed females ran 32% fewer laps than control females. Vape-exposed males showed a smaller reduction of 15% that did not reach statistical significance. The reduction was 10,025 laps per day larger in females than in males (p = 0.02). In females, the difference was present from the first weeks of exposure and was larger by the end of the study. The results demonstrate that vape exposure is associated with reduced voluntary aerobic activity. The analysis illustrates how nonlinear longitudinal modeling clarifies both the size and the temporal character of treatment effect in this experimental model. These findings reveal that nicotine-free components of vape aerosol reduce voluntary aerobic activity, highlighting an overlooked facet of vaping toxicity research which is particularly relevant for young and physically active users.

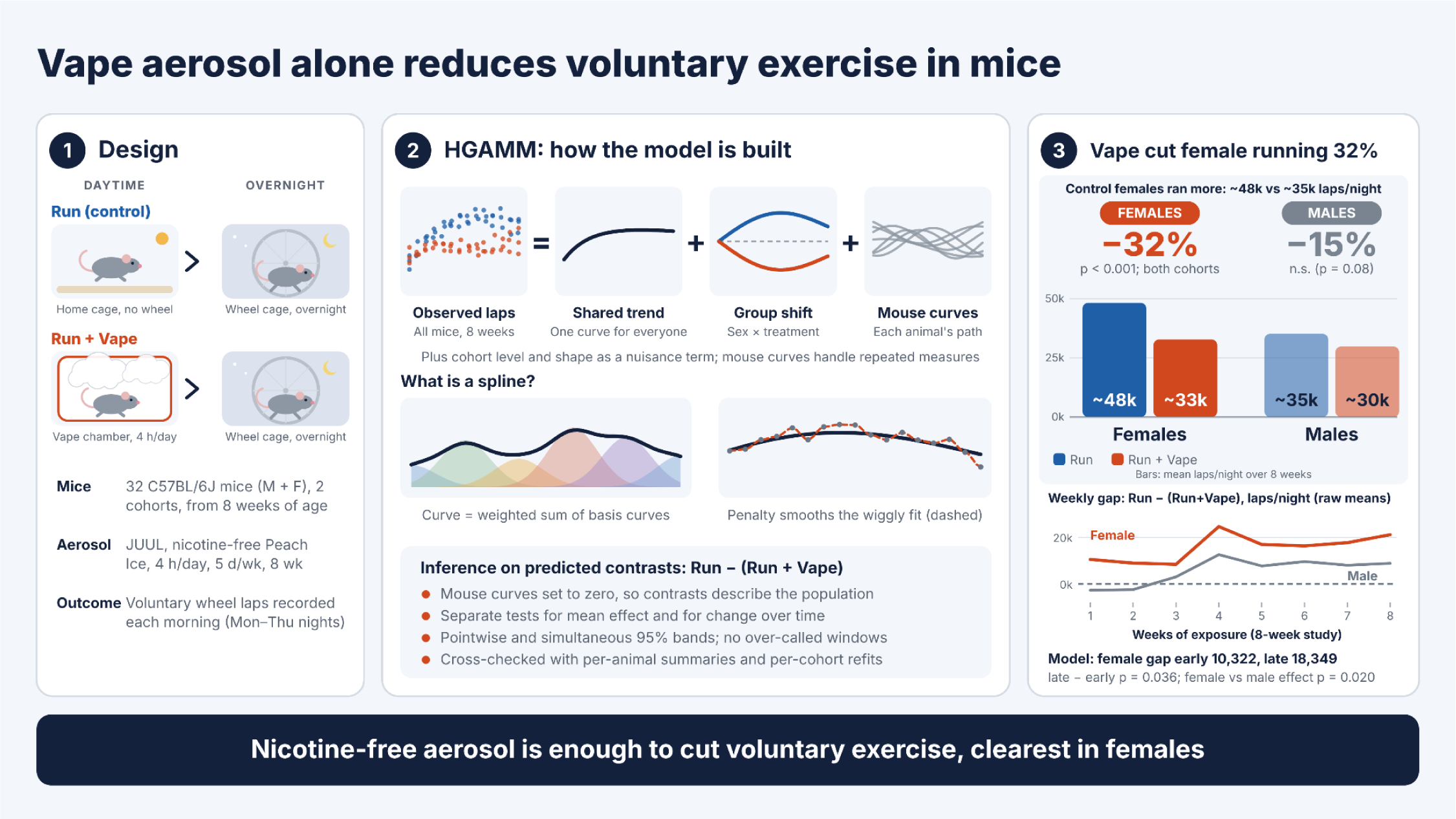

## 1. Introduction

Since vaping is seen as less harmful than smoking, many people adopt e-cigarettes as part of a healthier lifestyle. Websites touting the virtues of switching from smoking to vaping abound on the internet with a specific emphasis on advantages for athletes and individuals who exercise to stay healthy (Draper 2018; Hurley 2019; Lower Extremity Review 2019; Modern Man 2019; AFI 2020; Entwistle 2021; Baumeister 2022; Norman 2022). A recent meta-analysis found no articles on the benefits of exercise for vaping cessation or improvement of VO2max and/or VO2peak while such studies do exist for traditional smoking (Darabseh et al. 2022). The impact of vaping upon exercise performance is often suggested to be relatively benign for existing smokers who switch completely (George et al. 2019). Such rationalization may encourage smokers, fitness conscious individuals, and young athletes to adopt e-cigarettes as a healthier choice for their active lifestyle. Vaping is associated with compulsive exercise in college students along with a preference for nicotine vape juice (Ganson et al. 2022). Reports examining the intersection between vaping and exercise often focus upon improved performance outcomes compared to cigarette users and provide compelling rationales in favor of e-cigarette use (Hurley 2019; AFI 2020; Entwistle 2021; Baumeister 2022; Norman 2022). However, young vapers exhibit significantly reduced cardiorespiratory fitness, lower chronotropic response, and impaired skeletal muscle oxygen utilization during exercise (Simovic et al. 2024; Faisal et al. 2026), and popular coverage has begun to reflect these concerns for athletes and gym-goers (Hilliard 2019; O’Reilly 2019).

Young adults and adolescents are a physically active demographic with an average of 60 minutes of daily moderate-intensity activity (Bull et al. 2020). Conventional wisdom holds that athletic participation by students would decrease e-cigarette use as previously shown for tobacco smoking, but a national survey of high school seniors found otherwise. Adolescent athletes engaged in non-contact sports are at a lower risk of cigarette smoking, but the same protective influence of participation was not found with respect to e-cigarette use (Veliz et al. 2017). Studies of vaping demographics reveal seemingly paradoxical associations between vaping and exercise in young adults and adolescents. Those who vape display a 34.3% increased likelihood of exercise engagement compared to individuals who do not vape (Tiralla and Leung 2025; Rajbhandari-Thapa et al. 2022). This prevalent intersection between young vapers and their exercise regimens may have significant ramifications for the desired goal of increased fitness. Exercise improves cardiovascular, muscular, and metabolic health, making it a powerful tool in disease risk prevention. Multiple findings support exercise as a major contributor to prevention of cardiovascular disease as well as improvement of diagnosed obesity, type 2 diabetes, multiple sclerosis, and various other diseases (Qiu et al. 2023). Yet the relationship between physical activity level and inhalation of vape aerosols remains poorly defined.

Experimental animal models provide valuable insights regarding detrimental consequences of vape aerosol exposure (Alasmari et al. 2019; Chen et al. 2019; Jia et al. 2020; Carll et al. 2022; Esquer et al. 2022; Fried et al. 2022; Kim et al. 2022; Nogueira et al. 2022; Kim et al. 2024; Savko et al. 2025). In particular, our group reported pathological consequences in mice subjected to long term vape aerosol exposure (Esquer et al. 2022). We used a similar approach with lower exposure, which left the mice free to run over a two-month window. Age consideration of the mice is a major focus of our experimental design. We exposed mice starting at 8 weeks (56 days) for 8 weeks, (age at conclusion equals 105–112 days) roughly equivalent to human ages 15 to 22 (The Jackson Laboratory 2023) or alternatively 18 to 26 years (Dutta and Sengupta 2016). Two additional noteworthy design elements for the present study to mitigate stress upon subjects include 1) aerosol exposure via inhalation with minimal restraint via whole body chamber, and 2) unsupervised physical activity with voluntary running wheel activity taking advantage of normal nocturnal mouse behavior rather than forced exercise. Voluntary wheel running is an established method for physiological conditioning in mice (Goh and Ladiges 2015; Manzanares et al. 2018; De Bono et al. 2006; Hastings et al. 2022). Distances covered per evening are remarkable with averages of approximately 3 to 12 miles (Manzanares et al. 2018).

Resultant readouts of running wheel activity represented as laps per day per mouse over two months were interpreted using a hierarchical generalized additive mixed model (HGAMM) to reveal population-based characteristics for running distance between vaped versus non-vaped controls. Vape aerosol exposure decreased voluntary running, most clearly in females.

## 2. Materials and Methods

### 2.1 Animals, exposure, and design

#### Animals and housing

Male and female C57BL/6J were purchased from JAX and housed under a 12-h light–dark cycle with automatic light control at an ambient temperature of 70–72 °F, with a rodent maintenance diet (Teklad Global 14% protein) and water available ad libitum. The experiments began at eight weeks of age and continued for an additional eight weeks, encompassing the mice’s adolescent/young adult life. Mice were randomly assigned to treatment conditions identified as “Run” or “Run+Vape” conditions. Animal protocols and experimental procedures were approved by the Institutional Animal Care and Use Committee at San Diego State University (No. 25-068).

#### E-cigarette Exposure

Vapor was delivered to mice in whole body exposure chamber (SCIREQ inExpose) via JUUL-based aerosolization of nicotine-free Peach Ice e-liquid (70% vegetable glycerin / 30% propylene glycol; ORGNX) using the exposure protocol as previously described (Esquer et al. 2022). Briefly, the exposure profile used 3 second puffs at 20 second intervals with a vapor intake rate of 1.8L/min and fresh air cycling of 2.5L/min during aerosolization and 1.5L/min to clear the chamber. The cycle was repeated for 4 hours per day, 5 days per week, throughout the 8-week cycle. The exposure protocol and parameters have been validated by human vaping topography and follow recommended standards of e-cigarette exposure in laboratory models (Flouris et al. 2013; Dautzenberg 2015; Alasmari et al. 2018; Lechasseur et al. 2019; Soulet et al. 2019; Voos et al. 2019; Rapp et al. 2020; Gupta et al. 2021).

#### Running measurement and study design

Mice were housed individually Monday through Thursday night in cages equipped with running wheels and magnetic sensors to measure the number of laps run. The voluntary running wheel used for our study is similar to published odometer-based systems (De Bono et al. 2006). Laps were recorded the following morning resulting in 32 recorded running days over the eight-week exposure period (4 days recorded per week x 8 weeks). Mice were housed with unlimited wheel access except for time spent in the vaping chamber.

The dataset contains two independent experimental replicates, composed of male and female mice in both treatment groups, with four mice in each combination of cohort, sex, and treatment. The full design included 32 mice and a maximum of 1,024 daily observations across 32 recorded running days. Observations were excluded using two equipment-malfunction criteria. First, 80 observations with zero recorded laps were excluded when visual inspection documented wheel or sensor damage or misalignment. Second, 21 nonzero observations below 5,000 laps were excluded as suspected equipment malfunctions. The 5,000-lap threshold was established before the present statistical analysis and was based on historical wheel-running data from our laboratory, in which measurements from normally functioning equipment rarely fell below 5,000 laps per night. After these exclusions, 923 of 1,024 scheduled observations remained for analysis.Treatment was coded “R” (Run) or “R+V” (Run+Vape), sex as Female or Male, and cohort 1 and 2. The primary contrast was defined as Run minus Run+Vape within each sex, so positive values indicate greater modeled activity in Run mice.

### 2.2 Statistical Modeling Framework and Rationale

Analyses used R with the mgcv framework for model fitting (Wood 2017) and the gratia package for derivative and difference-curve estimation (Simpson 2026), with a Gaussian hierarchical generalized additive mixed models (HGAMM) fit by restricted maximum likelihood (REML) (Wood 2011). Gaussian models were retained as primary because they estimate treatment effects directly on the interpretable lap scale. Four other response families were fit using cross-validation (Section 3.5) and none improved predictive performance.

Since vape treatment was the effect of interest, differences in experimental replicates were accounted for, but were not of scientific interest and thus were defined as a fixed nuisance dimension (Pedersen et al. 2019). Group-level temporal patterns for the sex-by-treatment groups were expressed as deviations from a shared population curve using sum-to-zero (“sz”) factor-deviation smooths, assuming a shared underlying pattern and penalized accordingly. The primary model was:

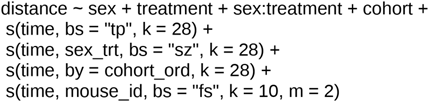

Here, distance is the number of wheel laps recorded for each mouse during an overnight measurement period; time is the first calendar day of exposure period, with measurements taken on days 2-5 of each 7-day week (e.g., days 2-5, 9-12); sex_trt is a four-level factor representing the combinations of sex and treatment; cohort_ord is cohort coded as an ordered factor; and mouse_id identifies individual mice. In the smooth terms, bs = “tp” specifies a thin-plate regression spline, bs = “sz” specifies sum-to-zero factor-smooth interactions for the four sex-by-treatment groups, and bs = “fs” specifies mouse-specific factor smooths. The parameter k sets the basis dimension for each smooth term, while m = 2 specifies the penalty applied to the mouse-specific smooth term.

Parametric terms estimate mean-level effects on the lap scale after accounting for nonlinear time (Wood 2003; Pedersen et al. 2019). The global thin-plate smooth estimates the shared trajectory; the sex-treatment deviation smooth allows the treatment trajectory to deviate from that shared curve while shrinking toward it; cohort terms absorb cohort-level differences in overall level and trajectory shape; and the mouse factor smooth captures each animal’s individual trajectory, handling the repeated-measures structure (Pedersen et al. 2019; Mundo et al. 2022; Sóskuthy 2017; Sóskuthy 2021). Double-penalty term selection (select = TRUE) allowed unsupported smooths to be shrunk toward zero.

### 2.3 Inference strategy

The treatment effect was decomposed into a time-averaged component and a time-varying component using tests that remain valid under penalized smoothing. Whether a mean-level treatment effect was present was assessed with a joint Wald test computed on the full linear predictor, with mouse terms set to zero, averaged over cohort and observed days with the unconditional coefficient covariance. Whether the treatment effect varied over time was assessed in two ways. The first used the sex-by-treatment factor-deviation smooth (its effective degrees of freedom) and the change in Akaike Information Criterion (AIC) when that smooth was added (Wood 2017). These AIC changes come from a three-step model progression fit by maximum likelihood: M0, no treatment; M1, adding the parametric treatment effect; M2, adding the treatment-by-time smooth. The second used early-versus-late contrasts estimated in a separate linear mixed model fit to the first two and last two recording weeks (days 2–5 and 9–12; days 44–47 and 51–54; 447 nights from 32 mice). That model included sex, treatment, period (early/late), their interactions, and cohort as fixed effects, with mouse-level random intercepts and random early/late shifts. Contrasts were averaged over cohorts with mouse terms excluded, using the unconditional covariance.

Time-averaged treatment contrasts (Run minus Run+Vape) were computed within each sex by averaging fitted contrasts across observed time points, using the model linear-predictor matrix and the model (unconditional) covariance matrix for Wald inference, with mouse-level terms excluded so that contrasts reflect population-level predictions. Pointwise treatment-difference curves across days were computed the same way (Marra and Wood 2012). Because the primary model contains no cohort-by-treatment interaction, the cohort term cancels in the Run minus Run+Vape difference. Time-averaged contrasts and difference curves are therefore identical across cohorts and are presented by sex. Difference curves and derivative curves (Section 2.4) were accompanied by simultaneous confidence bands obtained by posterior simulation from the coefficient covariance (Marra and Wood 2012; Simpson 2018), in addition to pointwise bands. All tests were two-sided with α = 0.05. P-values for the sex-specific, sex-difference, and time-course contrasts were not adjusted for multiple comparisons and should be read as exploratory.

### 2.4 Supporting Analyses

#### Rate of change

To identify periods when running increased or declined, we computed the first derivative of the fitted trajectory for each sex-by-treatment group on a 200-point grid (mouse terms excluded, averaged over cohort) by finite differencing the linear-predictor matrix, with simultaneous bands (Simpson 2014, 2017, 2018). Regions whose confidence band excludes zero were interpreted as periods of increase or decline.

#### Replication across cohorts

As a replication check, the model was fitted separately within each cohort, and the sex-specific treatment contrasts and confidence intervals were compared across cohorts.

#### Per-animal robustness analysis

Each mouse was reduced to summary quantities (total laps, mean laps, maximum single-day lap count, late-study slope) and analyzed with ordinary linear models, testing whether a simple per-animal summary reproduces the HGAM contrast.

#### Statistical resolution

For the sex-by-treatment interaction we report the minimum detectable effect at 80% and 90% power (two-sided α = 0.05) from the fitted standard error, framing the observed difference against the difference the study could resolve reliably.

#### Variance structure and diagnostics

We report the intraclass correlation (between-mouse variance as a fraction of between-mouse plus within-mouse variance), concurvity among the time smooths, basis adequacy (k.check), residual autocorrelation of within-mouse residuals, and the standard gam.check diagnostics.

#### Pooling and the interaction

The pooled primary model assumes treatment does not interact with cohorts. We tested this assumption three ways. First, we compared the pooled model to an interaction model by AIC and likelihood ratio test. The interaction model added a cohort-by-treatment level term and replaced the cohort shape smooth with a separate shape-deviation smooth for each cohort-by-treatment cell (reference: cohort 1, Run). Second, we tested for a cohort-by-treatment interaction in five per-mouse summaries using ordinary linear models: total laps, mean laps, peak laps, area under the running curve, and day of peak running. For each summary, we report the treatment-by-cohort coefficient from a model with sex, treatment, cohort, and treatment by cohort. We also report a joint F-test that drops all treatment-by-cohort terms from the full sex-by-treatment-by-cohort model. Third, we compared the pooled and interaction models by leave-one-mouse-out cross-validation, scoring each held-out mouse by root mean squared error and Gaussian log predictive density (LPD). Each held-out mouse was predicted at the population level, with predictive variance combining residual, between-mouse, and fit uncertainty. These fits used bam (fREML, discrete = TRUE) for speed. Effect sizes use REML with double-penalty selection. In-sample model comparisons (AIC, likelihood ratio test) use maximum likelihood without selection.

## 3. Results

### 3.1 Vape aerosol exposure reduced voluntary running in female mice

Running activity varied across time, sex, treatment, individual animal, and cohort, supporting a hierarchical longitudinal analysis rather than a cross-sectional or terminal-endpoint comparison (Figure 1). Averaged over the study, female Run mice ran substantially more than female Run+Vape mice: an estimated time-averaged difference of 15,444 laps per day (95% CI 9,424–21,464; p < 0.001), corresponding to a 32.0% reduction relative to the female Run mean. The male differential was smaller and not statistically resolved: 5,419 laps per day (95% CI −603–11,440; p = 0.08; estimated 15.5% reduction) (Figure 1B,C; Figure 2A,B).

**Figure 1.**
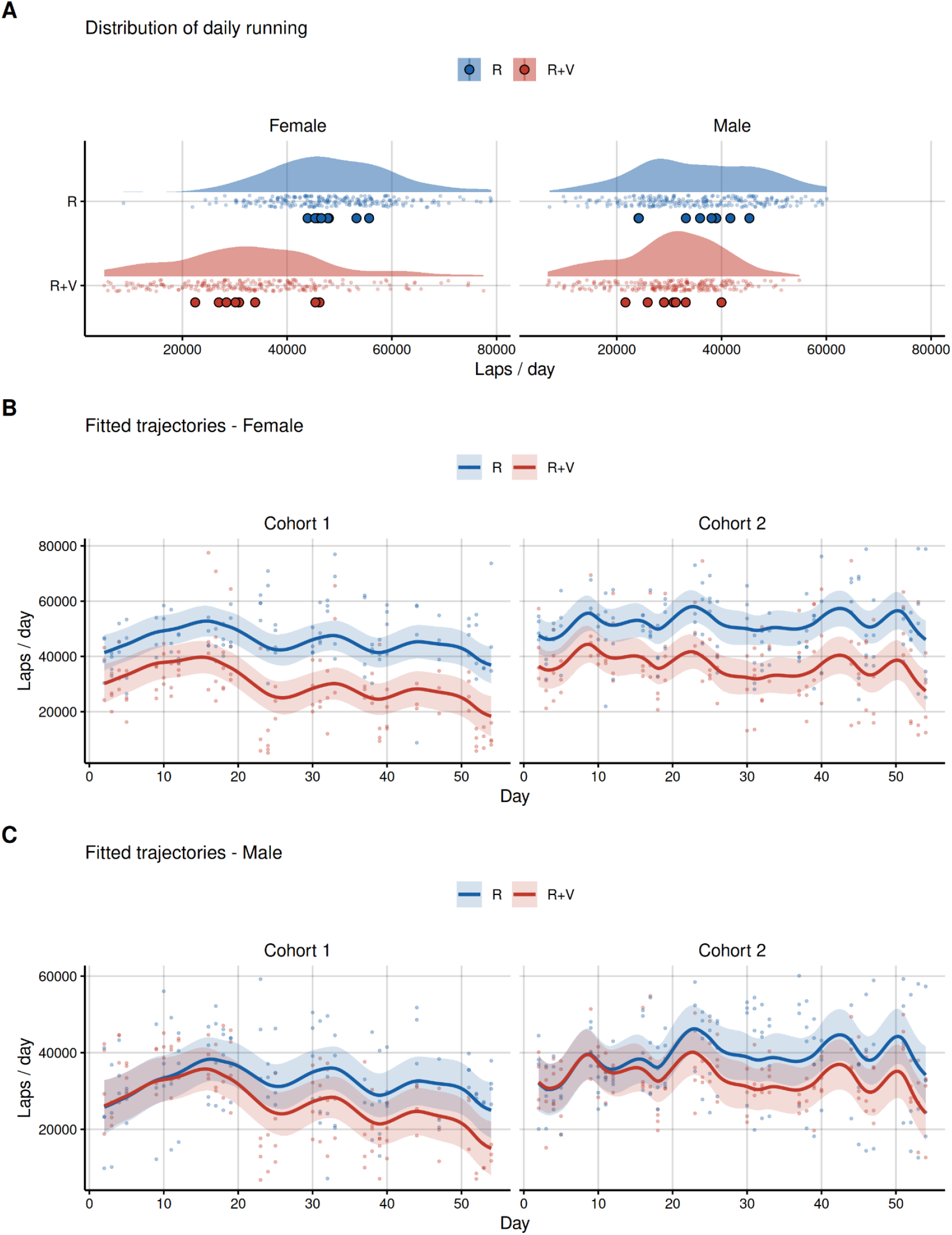
Nicotine-free vape aerosol exposure reduced voluntary running in females. The confidence interval for the smaller estimated reduction in males included zero. (A) Distribution of daily running by sex and treatment (horizontal raincloud: density of daily values with each mouse’s mean overlaid). (B, C) Fitted population trajectories (mouse effects excluded) with 95% confidence ribbons, faceted by cohort, for female (B) and male (C) mice where points are observed daily values. Female Run mice sit above female R+V mice across both cohorts while male trajectories overlap.

**Figure 2.**
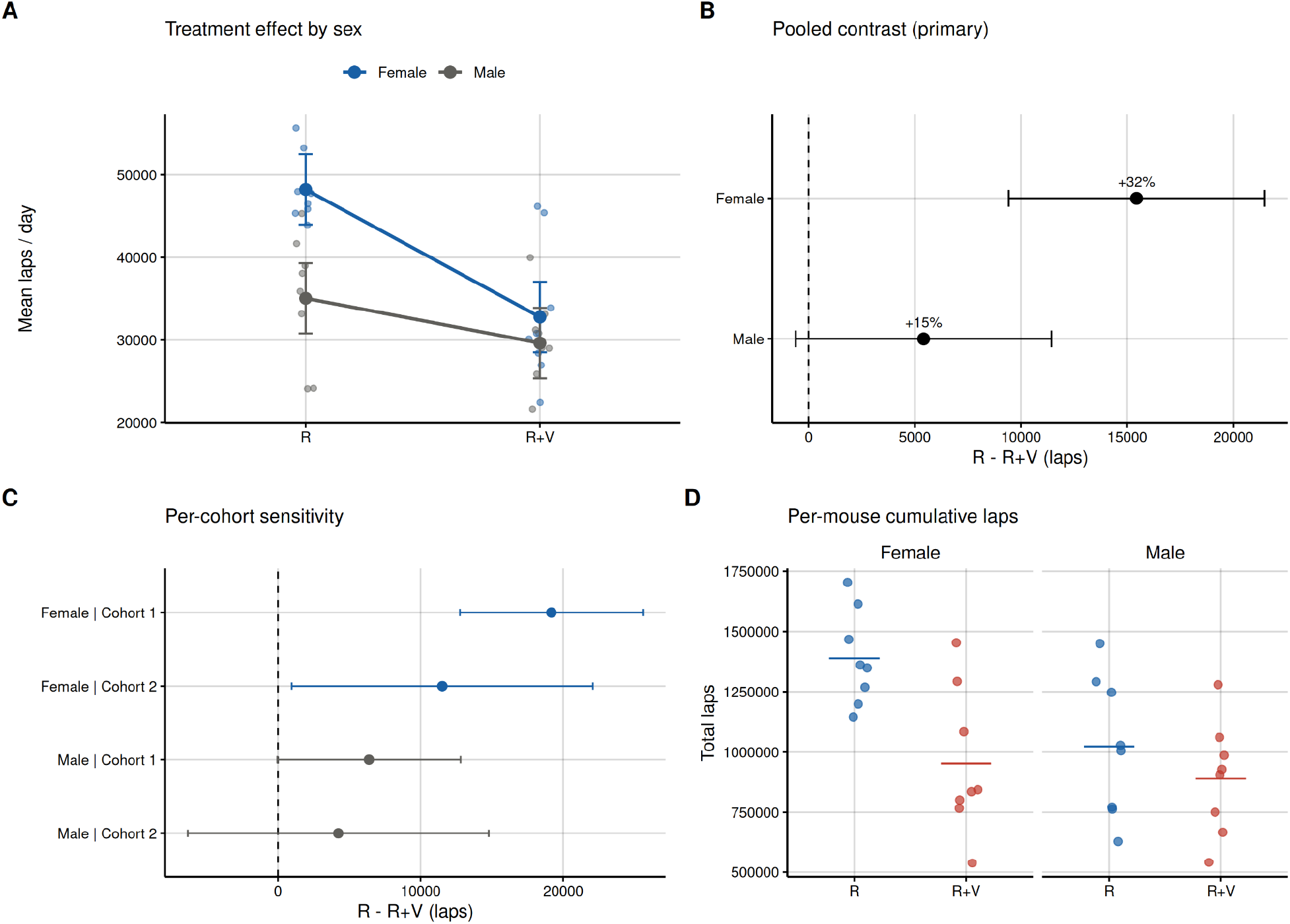
The female running deficit was consistent across cohorts and analyses. (A) Model-based mean laps for Run and Run+Vape connected within each sex; the difference in slopes corresponds to the sex-by-treatment interaction, with per-mouse means in grey and 95% CIs on the group means. (B) Pooled time-averaged contrasts (Run − [Run+Vape]) by sex with positive values indicating greater running in Run mice. Since the primary model contains no cohort-by-treatment interaction, contrasts are identical across cohorts and are shown once per sex. (C) Contrasts from models fit separately within each cohort. The female effect appears in both replicates. The male effect does not reach significance in either per-cohort refit (cohort 1 p = 0.052), although the cohort-1 male contrast is significant in the interaction model (Supplemental Table S3). (D) Per-mouse cumulative laps by group; an unconstrained summary reproduces the female effect.

The vertical R - (R+V) gap is constant across cohorts by construction, because the primary model contains no cohort-by-treatment interaction; cohorts differ in overall level and shape, not in the average magnitude of the treatment gap (Section 3.2).

The female effect also appeared in simple per-mouse summaries analyzed with ordinary linear models. Mean daily laps reproduced the female contrast (15,250 laps, p < 0.001) but not a male one (5,085 laps, p = 0.10). Total laps gave the same pattern (female 436,636 laps, p < 0.001; male 133,301 laps, p = 0.15; Figure 2D). Peak single-day laps (female 7,667, p = 0.081; male 6,451, p = 0.14) and the slope over the second half of the study (female p = 0.51; male p = 0.60) did not differ by treatment in either sex.

### 3.2 The reduction replicates in an independent cohort

The female reduction appeared in both replicates, fit separately. In cohort 1, there was a reduction by 19,172 laps (p < 0.001) and in cohort 2, it was 11,528 laps (p = 0.032). The estimated reduction in males was smaller, and its confidence interval included zero in the first cohort (6,383, p = 0.052) and the second (4,231, p = 0.43) (Figure 2C). The two female point estimates differ numerically, but their confidence intervals overlap substantially.

Formally testing whether treatment interacts with cohorts, the cohort-by-treatment interaction was preferred over the pooled model (LRT p < 0.001, ΔAIC ≈ 59).The parametric cohort-by-treatment coefficient was not significant (5,233, SE 4,240, p = 0.22). The two tests that do not depend on the collinear time smooths found no interaction in the size of the treatment effect. Per-animal summaries showed no interaction in total, mean, peak, or integrated running (all p > 0.27). Day of peak running did differ (treatment-by-cohort estimate 29.8 days, p = 0.002), indicating the cohorts differed in when running peaked rather than in how much the treatment reduced it. In leave-one-mouse-out cross-validation, the two models predicted held-out animals no better than one another, and the apparent in-sample gain tracked cohort rather than treatment (Supplemental Table S1). As the experimental question under investigation here concerns whether vaping affects running, and because the evidence of a cohort-specific treatment effect was not consistent across analyses,the pooled model was retained as the primary summary. The model comparison is in Supplemental Table S1, and the interaction model’s cohort-specific contrasts are in Supplemental Table S3 and Supplemental Figure 2. In that model, the male contrast was significant in cohort 1 (8,384 laps, p = 0.025) but not in cohort 2 (2,887 laps, p = 0.44).

### 3.3 The estimated deficit is present early and larger later

The treatment effect in females was not a constant offset. A joint Wald test on the full linear predictor confirmed a mean-level effect (χ²(2) = 28.39, p < 0.001). Adding the treatment terms improved AIC by only 0.38 (M0 to M1). Without treatment terms, the mouse-specific smooths absorb the group difference as per-mouse offsets, so moving it into the fixed effects barely changes the fit (model EDF fell from 109.2 to 108.1). AIC therefore cannot detect a between-group effect in this design, and inference on the mean effect rests on the Wald test. The sex/treatment interaction smooth carried appreciable structure (effective degrees of freedom ≈ 4.6), and adding it improved AIC only marginally (ΔAIC ≈ 2.7, M1 to M2). In the window model, the female deficit was present in the first two weeks (weeks 1–2: 9,652 laps, 95% CI 1,177–18,127; p = 0.026) and was larger by the end of the study (weeks 7–8: 20,474 laps, 95% CI 11,931–29,016; p < 0.001). A late-versus-early contrast supported the increase (10,821 laps, 95% CI 1,081–20,562; p = 0.029). In males, the early difference was not resolved (−2,637 laps, 95% CI −11,064 to 5,789; p = 0.54) and the late difference was imprecise (7,244 laps, 95% CI −1,345 to 15,834; p = 0.098). The male late-versus-early change was nominally positive (9,882 laps, 95% CI 149–19,614; p = 0.047) but did not differ from the female change (difference 940 laps, 95% CI −12,826 to 14,706; p = 0.89), so the time course does not distinguish the sexes at this sample size. No group trajectory showed a sustained period of change. Simultaneous bands for the first derivative excluded zero only briefly, at 3 of 200 grid points in female Run mice. The female treatment-difference curve was positive across the study, while the male difference curve straddled zero throughout (Figure 4A,B). The vaping-associated reduction in female running is therefore best described as present early and larger late, rather than as a steadily widening gap. The derivative analysis describes each group’s trajectory and does not test the gap directly.

One feature of this design constrains how the temporal result should be interpreted. The population time smooths were highly collinear (concurvity approaching 1; Figure 3B), meaning the model could not separate the effects of the global, cohort, and sex-treatment time-dependent terms. Therefore, individual effective degrees of freedom or nominal p-values of the time smooths are not interpreted.

**Figure 3.**
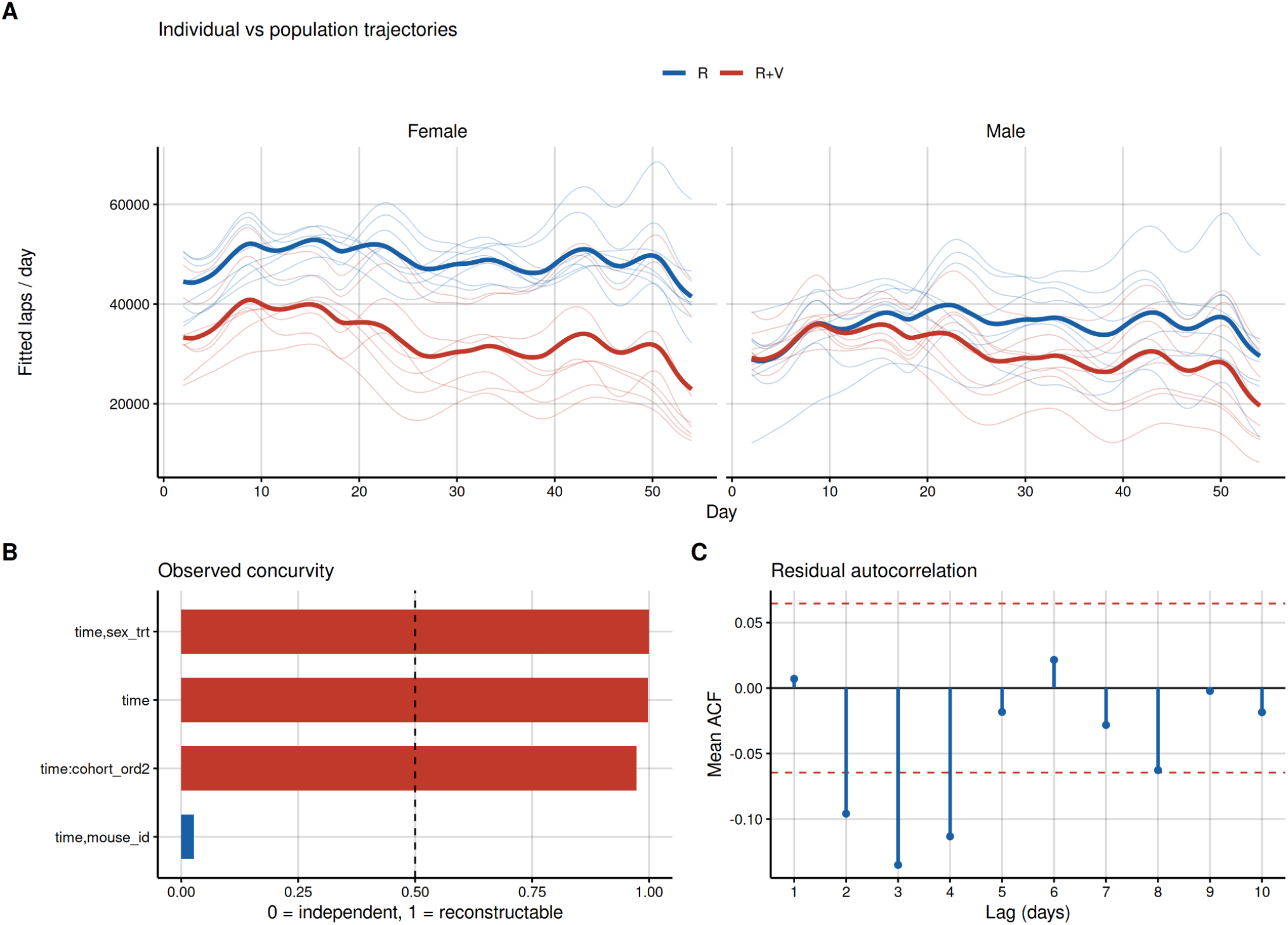
Individual trajectories differed by treatment in females but showed greater overlap in males. (A) Individual fitted trajectories for all 32 mice (faint) behind the population trajectories (bold), by sex. Run trajectories lie above Run+Vape trajectories in females and overlap in males. (B) Population time smooths are highly collinear (values near 1) while the mouse factor smooth is well identified (≈ 0.03), justifying inference by contrasts rather than by individual smooth significance. (C) Mean within-mouse residual autocorrelation by lag, in recorded nights. Lag-1 autocorrelation is near zero. Lags 2–4 show small negative values (about −0.10 to −0.13) that cross the approximate 95% bounds. Negative autocorrelation of this size is expected after fitting a smooth trajectory to each mouse and would make standard errors conservative rather than too small.

This collinearity does not undermine the mean-effect and time-varying conclusions above, which rest on prediction based contrasts, simultaneous difference bands, and the derivative analysis, none of which rely on attributing structure to a single smooth. It also explains why the interaction model was favored in Section 3.2. The interaction term added more flexibility to the model and thus can absorb structure already captured by the cohort shape smooth, but cannot localize it to a single factor. The mouse factor smooth showed low observed concurvity (≈ 0.03), indicating it captured independent, within-animal variation.

### 3.4 Evidence of a sex difference in the treatment effect, with limited precision

The estimated reduction in females was 10,025 laps greater than that in males (95% CI 1,513–18,538; p = 0.021). The investigation was able to reliably identify between-sex differences of 12,168 laps at 80% power and 14,078 at 90% with only four mice per cell. The wide confidence interval indicates that the magnitude of the observed sex difference was estimated with limited precision. (Figure 4C). As a result, the sex difference is supported but modest, its magnitude is coarsely resolved, and it is not corrected for the several reported contrasts. The male effect was smaller and not significant.

**Figure 4:**
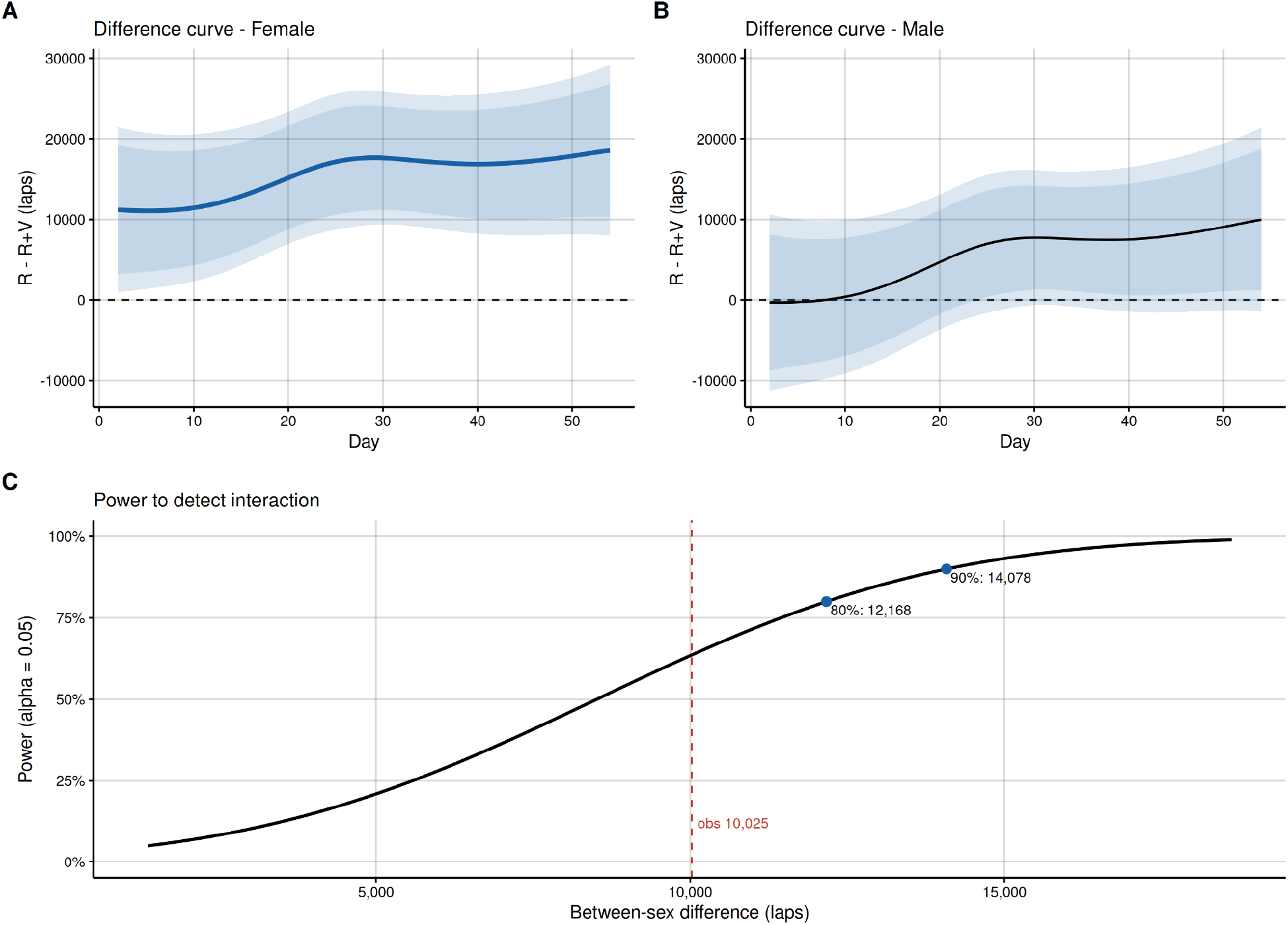
Evidence of a sex difference, with limited precision. (A, B) Treatment-difference curves (Run − [Run+Vape]) for female (A) and male (B) mice are shown on a common y-axis over the course of the study with pointwise and simultaneous 95% bands. Days where zero is not included in the simultaneous confidence band are shown by bold segments. The male difference straddles zero throughout, while the female difference rises from ∼11,000 laps in weeks 1–2 to ∼18,500 in weeks 7–8. (C) The power to identify the sex-by-treatment difference as a function of its actual magnitude, with the lowest detectable impact shown at 80% (12,168 laps) and 90% (14,078). The observed difference of 10,025 laps and its 95% CI of 1,513–18,538 are shown for reference.

### 3.5 Model adequacy

Between-mouse differences accounted for roughly a quarter of total residual variation (intraclass correlation ≈ 0.27), indicating substantial individual differences and supporting the per-mouse terms. The effective degrees of freedom provide a measure of model flexibility, with the per-animal smooths (≈ 70.0) and cohort trajectory-shape difference (≈ 13.7) being the most flexible, followed by global trend at ≈ 9.9 and the sex-by-treatment deviation at ≈ 4.6. The individual variability is visible directly in the per-mouse trajectories (Figure 3A). The k-index was ≈ 0.93 (p ≈ 0.01–0.02), but effective degrees of freedom were well below the basis limits. Mean within-mouse lag-1 residual autocorrelation was near zero (≈ 0.006; Figure 3C). Small negative autocorrelation appeared at lags 2–4 (about −0.10 to −0.13), consistent with fitting a smooth to each mouse’s short series, and would make standard errors conservative. Standard diagnostics (Supplemental Figure 1) showed approximately linear Q–Q behavior through the central distribution with modest tail deviation, a symmetric residual histogram, and strong observed-versus-fitted association, with mild heteroscedasticity typical of biological activity data. The model explained 65% of the deviance. Because overall fit is dominated by individual variation, inference on treatment is based on the population-level contrasts and tests described above rather than on overall model fit.

Animal-by-animal scores against the Gaussian baseline were used to assess four additional response families based on how well each predicted held-out mice (Supplemental Table S2; negative values imply worse prediction than Gaussian). There was no benefit to modeling heavy tails or non-constant variance: Gaussian location-scale ΔLPD +0.001 (SE 0.022, p = 0.96; RMSE 11,510 vs. 11,537 for Gaussian, favored in 20/32 mice), scaled-t 0.000 (p = 0.95). Negative binomial was poorer by LPD (−0.055, p = 0.03), with a similar but not resolved RMSE difference (+243 laps, p = 0.063), whereas Tweedie was no better (−0.007, p = 0.59) and less accurate (RMSE p = 0.007). None of the alternative families resolvably improved out-of-sample predictive performance. Thus, Gaussian was kept, maintaining effects on the interpretable lap scale.

## 4. Discussion

Long term consequences of e-cigarette use (vaping) among adolescents and young adults remain poorly characterized. In 2024, 7.1% of high school and 2.6% of middle school students reported vaping in the past 30 days, revealing the habit often begins at an early age (Jamal et al. 2024; Sun et al. 2026). E-cigarette users on average performed at a lower fitness level than non-users in a survey of active duty US army soldiers (Dinkeloo et al. 2020). Vaping is a serious concern because of the potential for compromised fitness in military ranks (Little et al. 2015; Chin et al. 2018; Hall et al. 2018; Dinkeloo et al. 2020; Lin et al. 2020; Colesar and McCollum 2021; Keltner et al. 2021; Gill et al. 2022) as well as for other groups engaged in physically demanding pursuits such as emergency workers, first responders, and athletes (Veliz et al. 2015; Veliz et al. 2017; Young et al. 2020; Gross 2021). Furthermore, increasingly sedentary lifestyles in our society have raised awareness of health risks and renewed urgency for voluntary exercise activities (Fiuza-Luces et al. 2018; Lavie et al. 2019; Channon 2020; Guo et al. 2020; Qiu et al. 2023).

Findings from this report demonstrate chronic vaping exposure is associated with reduced voluntary aerobic activity most clearly in female mice. The female reduction was large (∼32%), appeared consistently across two cohorts, survived independent per-animal analysis (per-mouse linear models and per-cohort refits), and was not changed by modeling cohort as an interaction. The convergence of model contrasts, per-mouse summaries, and cohort-wise refits gives the female result strong internal support for a study of this size.

Running was highly nonlinear over the course of the eight weeks, and this nonlinearity was a key component of the treatment effect. In females, the gap began early and was estimated to be larger by the end of the study. Characterizing it that way, rather than as a simple decline or a stable offset, required modeling the nonlinear time course directly.

The apparent sex difference should not be over-interpreted mechanistically. The difference between sexes was statistically supported yet imprecisely estimated, so it is best regarded as provisional. Possible contributors to a genuine sex difference are pulmonary or cardiovascular responses, inflammatory or hormonal signaling, or differences in running motivation which cannot be distinguished here, as the study quantifies behavior, not mechanism. Sex-related differences in voluntary wheel-running activity are themselves well documented in mice (Bartling et al. 2017).

The e-liquid used here contained no nicotine, so the observed reduction in voluntary running cannot be attributed to nicotine and instead reflects effects of the inhaled aerosol vehicle (vegetable glycerin (VG) and propylene glycol (PG)) and flavorant alone. The non-nicotine component of vaping exposure is often overlooked in a nicotine-focused literature, but the aerosol itself is sufficient to depress voluntary aerobic activity in females in this model. Most of the epidemiology cited in the Introduction concerns nicotine products, so it applies to these results only indirectly. The present findings speak to the vehicle-and-flavor aerosol, and a nicotine-containing comparison would be a natural next step.

This result sits within a small but informative body of rodent work testing whether the vehicle alone, without nicotine, is sufficient to impair exercise. The most direct precedent is a study in which nicotine-containing e-cigarette aerosol reduced treadmill running speed in male mice by 8%, but the vehicle (PG/VG) delivered without nicotine reduced running speed by an even larger 11% and impaired recovery from muscle injury, indicating that the vehicle alone can measurably impair exercise performance and is not a physiologically inert carrier (Nogueira et al. 2022). This follows other studies showing that vegetable glycerin and propylene glycol aerosols induce airway inflammation and disrupt normal ion-channel function in human airway cells and large-animal models (Kim et al. 2022; Kim et al. 2024), which suggests a possible respiratory path where vehicle itself could reduce the capacity or willingness of an animal to run. A separate study of female mice with acute exposure to e-cigarette vapor found that zero-nicotine vapor exposure showed a small non-significant trend of reduced swimming endurance and grip strength, with much more prominent impaired physical capabilities emerging when exposed to nicotine-containing aerosols (Chen et al. 2019). Our findings are more consistent with Nogueira et al. (2022). Vehicle and flavorant can be sufficient on their own, and several design differences may help explain the discrepancy. Our exposure was chronic (eight weeks of whole-body inhalation exposure, five days per week) rather than acute (14 days of brief chamber dosing), our outcome was voluntary wheel running rather than forced swimming or grip strength, and the two studies used different e-liquid formulations and delivery devices. Whether non-nicotine vaping constituents can independently impair voluntary aerobic behavior may therefore depend on exposure duration and the specific behavioral assay used, an open question this study cannot fully resolve but to which it adds a chronic, voluntary-exercise data point.

The analysis illustrates several practices that generalize to preclinical longitudinal studies. Nuisance factors such as experimental batch should be adjusted for, but the form of the adjustment must match the number of levels. With few levels, variance estimation would be unreliable, so a fixed effect is preferable. By separating the time-varying test from the mean-effect test, it is possible to distinguish between “is there an effect” and “does it change over time,” both of which need inference that is still valid after penalized smoothing. Pairing pointwise with simultaneous bands prevents over-declaring localized significance and confirming a hierarchical result with a simple per-animal summary guards against the suspicion that an effect is an artifact of the smoothing.

### 4.1 Limitations

The study tested one nicotine-free flavored formulation and cannot identify the contribution of its individual components. Each cohort-by-sex-by-treatment cell contained four mice, limiting power to detect smaller effects. Thus, although the sex-by-treatment difference was detected, the study was powered to resolve its magnitude only coarsely. The pre-exposure baseline period was not measured. Because exposure was in effect throughout the recorded window, we cannot directly rule out pre-existing group differences, although random assignment to condition mitigates this concern. Population time smooths were highly collinear in this two-cohort design, so time-varying structure cannot be localized to a specific smooth. We therefore base inference on contrasts and nested tests, which are defined on predictions and remain valid under this collinearity (Section 3.3), rather than on individual smooth significance. Gaussian models showed mild heteroscedasticity and tail deviation, common in activity data; location-scale and heavy-tailed alternatives offered no predictive improvement. Finally, model-based difference and derivative curves identify modeled regions of change but do not establish a precise biological onset.

## 5. Conclusion

Chronic vaping exposure was associated with a large, reproducible reduction in voluntary running in female mice while the estimated reduction in males was smaller, and its confidence interval included zero. The deficit was present from the first weeks of exposure and estimated to be larger by the end of the study, although no sustained period of change was identified. The effect was observed in both experimental cohorts of animals and was robust across the longitudinal and per-animal analyses. Because the e-liquid contained no nicotine, the reduction reflects the vehicle and flavorant found in the ORGNX nicotine-free peach ice formulation. Vape aerosol alone is sufficient to reduce voluntary exercise in female mice. That cost falls precisely on young, active users who are often reassured that vaping is the safer choice.

## Supporting information

All Supplemental Data

## AI Disclosure Statement

The statistical methodology, theoretical results and interpretation in this article were developed by the authors without AI assistance. Claude (Anthropic) was used to aid in the generation of the R code and graphical abstract supporting the manuscript. All AI-generated code was reviewed by the authors, and its implementation choices were checked against the methodology. Claude was used solely for grammatical revision and language refinement to improve the clarity, readability, and fluency of the text. The original draft was written entirely by the authors who retain full responsibility for the content, interpretation, and final version of the manuscript.

## Data Availability

All code and running data is available at https://github.com/ddawood27/Vape-Aerosol-Exposure-Decreases-Exercise-Performance-in-Mice-R-Scripts.

## Acknowledgements

The authors extend their deepest appreciation to members of the Sussman Laboratory, who provide invaluable assistance and expertise in developing vaping-related studies and information.

---

**Table 1:** Treatment-effect tests. The linear predictor is used to calculate all contrasts (Section 2.3). The AIC changed by ≈ 2.7 (smooth edf = 4.6) when the sex-by-treatment time smooth was included; its nominal p-value is not utilized since it is unreliable under the observed concurvity. Due to the non-integer effective degrees of freedom of these models, a sequential likelihood-ratio test is not employed.

| Question | Basis | Result |
| --- | --- | --- |
| Mean treatment effect? | Joint Wald that the time-averaged R – R+V contrast is zero in both sexes | $\chi^2(2) = 28.39, p < 0.001$ |
| Sex difference in effect? | Female – Male contrast | 10,025 laps, SE 4,343, $p = 0.021$ |
| Does the effect change over time? | Late (weeks 7-8) minus early (weeks 1-2) female contrast | 10,821 laps, $p = 0.029$ |
| Any supported rate of change? | First derivative of each sex-by-treatment trajectory, simultaneous bands | Band excluded zero at 3 of 200 grid points ($\approx 1$ day, female Run); none in other groups |

**Table 2:** Time-averaged treatment contrasts (Run − Run+Vape), by sex. All three rows are prediction-based averages of the fitted difference across observed days, averaged over cohort, with mouse terms excluded.

| Sex | Contrast (laps) | 95% CI (laps) | % Reduction | P-value |
| --- | --- | --- | --- | --- |
| Female | 15,444 | 9,424 – 21,464 | 32.0% | <0.001 |
| Male | 5,419 | –603 – 11,440 | 15.5% | 0.08 |
| Female - Male Interaction | 10,025 | 1,513 – 18,538 | – | 0.021 |

