## Supplementary material for "Vape Aerosol Exposure Decreases Voluntary Exercise in Mice": All Supplemental Data

### Supplement

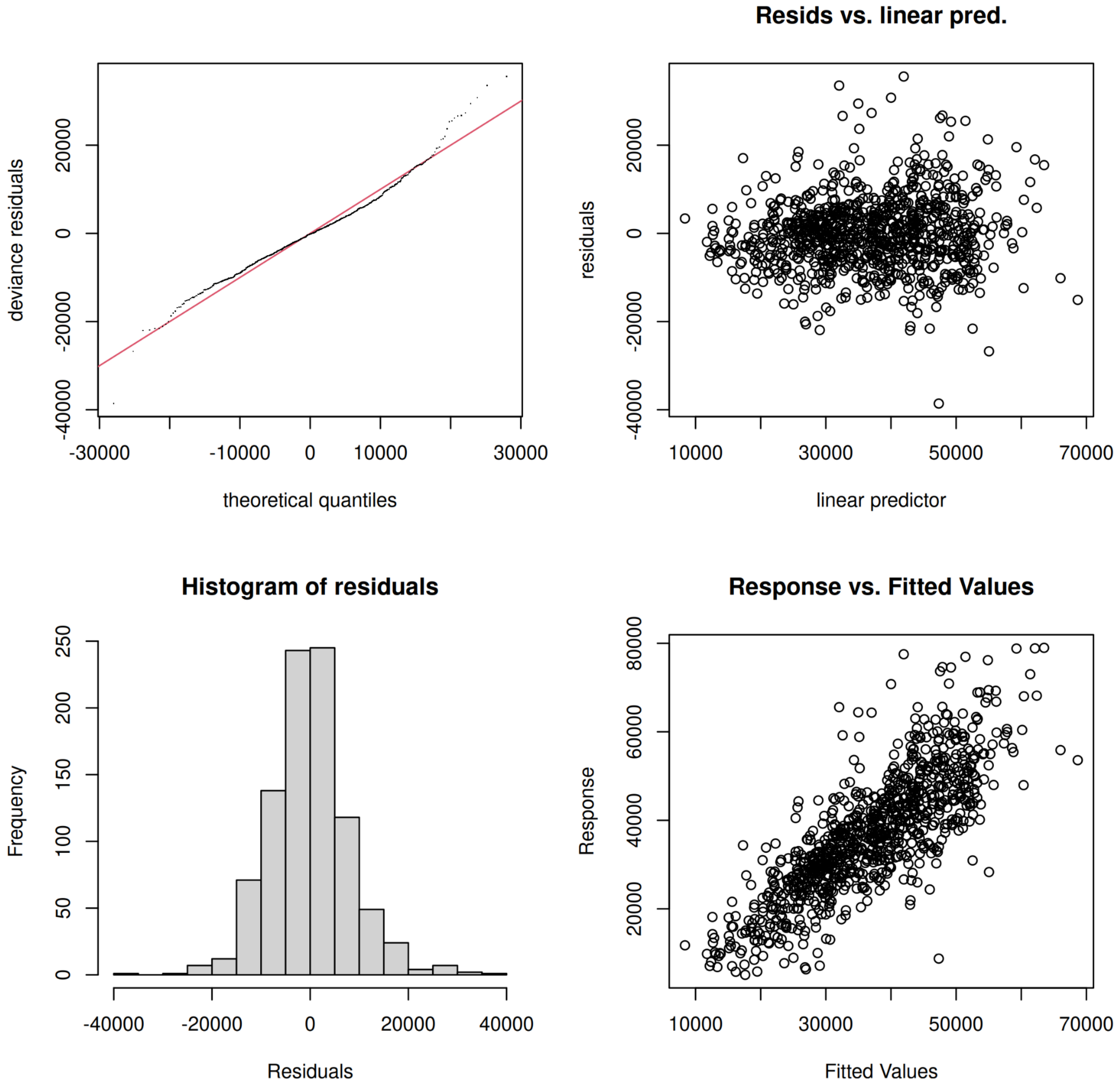


**Supplemental Figure 1. Model diagnostics.** Residual Q–Q plot, residuals versus linear predictor, residual histogram, and observed versus fitted values, supporting the Gaussian model for lap-scale inference while indicating mild tail deviation and heteroscedasticity typical of running-wheel data.

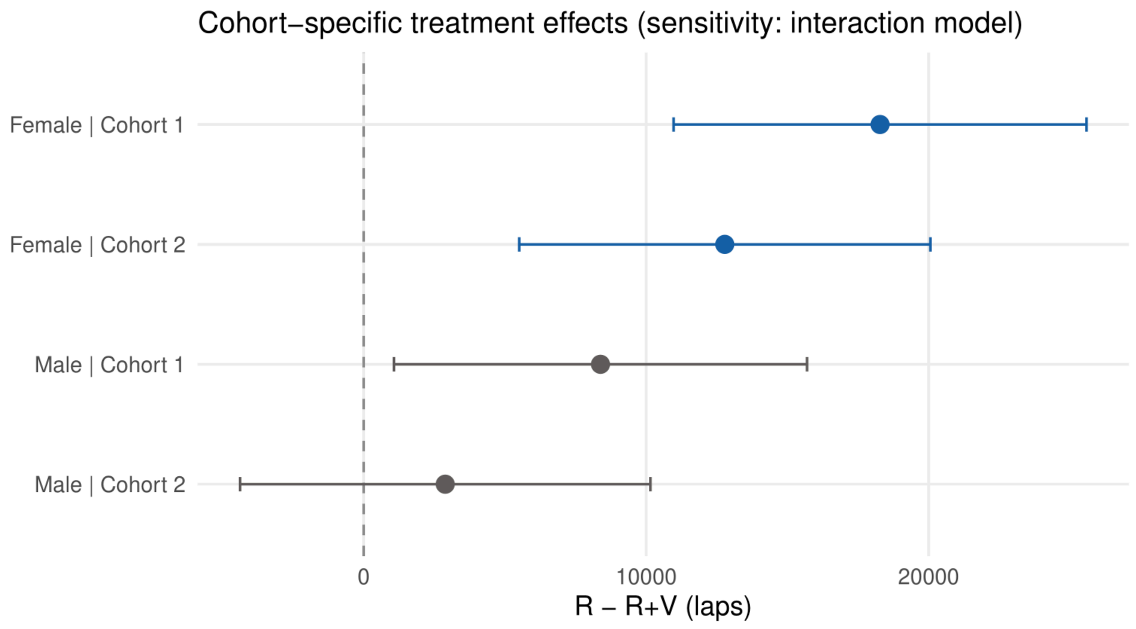


**Supplemental Figure 2. Cohort-specific treatment effects in the interaction model.** Run − Run+Vape contrasts with 95% CIs for each sex and cohort, from the sensitivity model that allows the treatment effect to differ by cohort (values in Supplemental Table S3). The dashed line marks zero.

| | Pooled (primary) | Cohort x Treatment |
|---|---|---|
| Total EDF | 101.4 | 108.1 |
| AIC (ML) | 19,451.6 | 19,393.0 |
| Parametric cohort × treatment coefficient | – | 5,233 (SE 4,240, $p = 0.22$) |
| Held-out ΔLPD vs pooled (95% CI) | – | +0.009 (−0.035 to +0.053); Wilcoxon $p = 0.30$ |
| Held-out ΔRMSE vs pooled (95% CI) | – | −135 laps (−551 to +281) |
| Mice better predicted | 11/32 | 21/32 |
| Per-cell ΔLPD: cohort 1 Run / Run+Vape | – | +0.033 / +0.052 |

| | | |
|---|---|---|
| Per-cell ΔLPD: cohort 2 Run / Run+Vape | – | −0.021 / −0.029 |

**Supplemental Table S1: Cohort × treatment interaction: supporting detail for Section 3.2.** Leave-one-mouse-out cross-validation, 32 refits per model, paired by animal. The interaction model's advantage tracks cohort rather than treatment arm, and its own cohort-by-treatment coefficient is not resolved. EDF, AIC, and LRT are from ML fits; the parametric coefficient is from the REML fit with double-penalty selection.AIC changed by 0.38 when the parametric treatment terms were added (M0 to M1). See Section 3.3 for why AIC is not informative for this step.

| Family | Mean RMSE | ΔLPD vs Gaussian (SE) | Paired t-test | Mice Favoring Alternative |
|---|---|---|---|---|
| Gaussian | 11,537 | – | – | – |
| GaussianLS | 11,510 | +0.001 (0.022) | 0.96 | 20/32 |
| Scaled-t | 11,570 | 0.000 (0.007) | 0.95 | 19/32 |
| Tweedie | 11,734 | −0.007 (0.013) | 0.59 | 13/32 |
| Negative Binomial | 11,780 | −0.055 (0.024) | 0.028 | 14/32 |

**Supplemental Table S2.**

**Response-family comparison, leave-one-mouse-out, paired against Gaussian.** Five families were scored on each held-out mouse by log predictive density (LPD) and RMSE (laps). ΔLPD is the mean per-mouse difference from Gaussian (positive favors the alternative); "mice favoring alternative" counts animals better predicted than by Gaussian. No alternative beat Gaussian by a supported margin, so Gaussian was retained.

| Sex | Cohort | Contrast, Run-Run+Vape (laps) | 95% CI (laps) | p-value |
|---|---|---|---|---|
| Female | 1 | 18,281 | 10,972 – 25,590 | < 0.001 |

| | | | | |
|---|---|---|---|---|
| Female | 2 | 12,784 | 5,505 – 20,063 | < 0.001 |
| Male | 1 | 8,384 | 1,072 – 15,696 | 0.025 |
| Male | 2 | 2,887 | −4,378 – 10,151 | 0.44 |

**Supplemental Table S3: Cohort-specific treatment contrasts from the interaction model (sensitivity analysis).** Time-averaged Run − Run+Vape contrasts from the cohort-by-treatment interaction model, fit by REML with double-penalty selection. Contrasts are averaged over observed days with mouse terms excluded, using the unconditional covariance. The effect is in the same direction in every cell. The male cohort-1 contrast reaches significance here but not in the per-cohort refit (6,383 laps, $p = 0.052$; Section 3.2).
